# Forecasting bryozoan assemblage dynamics under simulated climate change

**DOI:** 10.1101/2025.10.31.685734

**Authors:** Mira Baer, Korinna T. Allhoff, David K. A. Barnes, Franziska Koch

## Abstract

The shallow Antarctic continental shelf experiences strong physical disturbances in the form of ice-scour which play a key role in maintaining biodiversity. Due to climate change, both the extent and duration of sea ice cover is expected to rapidly decline, leading to complex shifts in disturbance regimes with unknown impacts on successional dynamics and biodiversity in benthic communities. We introduce a simulation model to study the assemblage dynamics of Bryozoa, which are sessile, suspension feeding animals and key pioneers in the shallows. The model captures colonisation events, colony growth as well as intra- and interspecific overgrowth competition for space between colonies. Mortality due to predation is modelled as the removal of zooids within small areas of the model world. Using the model, we simulate various disturbance regimes, e.g. by varying the timing of ice-scour events, the length of the growing season and the spatial distribution of predation events. We find that the timing of ice-scour events throughout the growing season has only minor short-term effects on successional dynamics in bryozoan assemblages, while an extended growing season substantially accelerates succession in the long term. We furthermore find that relatively rare but large predation events lead to a slower recovery, whereas relatively frequent but small events result in a faster succession with higher overall abundances. These results highlight that in order to understand how benthic biodiversity will be impacted by climate change, it is necessary to consider the interplay between biotic interactions and complex changes in physical disturbance regimes.

**Highlights:**

- We use an individual-based model to simulate climate change impacts on bryozoan assemblages.
- Warming leads to complex shifts in timing and spatial distribution of disturbances.
- Successional dynamics are affected by changes in growing season and predation events.
- Interplay of biotic interactions and physical disturbances drives benthic biodiversity.

## 1 Introduction

The shallow Antarctic shelf is considered one of the most naturally disturbed areas on earth [Barnes and Conlan, 2007, Gutt, 2001, Gutt and Starmans, 2001]. Icebergs regularly collide with the seabed and kill most organisms on their way, a process termed ice-scouring [Barnes et al., 2024a, Brown et al., 2004]. Despite these catastrophic disturbances, the Antarctic Benthos is very species rich [Arntz et al., 1994, Clarke and Johnston, 2003, Griffiths et al., 2024]. Ice-scours leave behind open space, which is available for recolonisation. The early stages of this succession are dominated by fast-growing pioneer species, but these are later replaced by slower-growing species that are better at competing for space [Barnes and Rothery, 1996, Zwerschke et al., 2021]. Without disturbance, these climax species would eventually completely exclude the pioneers. Frequent disturbance by ice-scouring that regularly resets the succession enables the coexistence of pioneers and climax species, and is thus key to maintaining the high biodiversity of the region [Brown et al., 2004, Barnes and Conlan, 2007, Robinson et al., 2021].

The Southern Ocean has remained at constant, cold temperatures for millions of years, but climate change is now causing multiple interacting physical changes [Pachauri et al., 2014], with strong impacts on benthic ecosystems. Most species are endemic and considered to be highly sensitive to warming [Peck et al., 2004, Barnes and Peck, 2008]. Additionally, dynamics are strongly linked to sea ice cover [Gutt, 2001]. The duration and extent of sea ice is already decreasing, with ice now thawing earlier and faster [Gilbert and Holmes, 2024, Cook et al., 2005, Parkinson, 2014]. Combined with glacier retreat and ice-shelf collapses, this is leading to an increase in ice-scouring rates [Barnes et al., 2024a], which is predicted to further continue within the next decades.

How increased ice-scour frequencies affects benthic communities has been studied by comparing community structure across locations with different disturbance frequencies. For example, [Robinson et al., 2021] used a depth gradient for this purpose, as shallower areas are hit more frequently than deeper ones. At intermediate depths and ice-scouring rates, species richness was found to be highest, while both very high and very low disturbance frequencies lowered species richness. These observations can be explained through the intermediate disturbance hypothesis [Connell, 1978, Wilkinson, 1999, Roxburgh et al., 2004, Miller et al., 2011, Sheil and Burslem, 2013, Moi et al., 2020]: If the succession is reset more often, later successional stages are never reached and the community consists only of fast-growing pioneers.

However, increased ice-scour frequency is only one of several potential ways in which climate change might affect the succession in Antarctic benthic systems. Rising tem peratures might also impact community dynamics by directly influencing growth rates [Peck et al., 2004] or by modifying biotic interactions like competition for space [Ashton et al., 2017, Barnes et al., 2021]. Such direct effects are already difficult to study in the field, but it is possible for example through the use of submerged heated panels that are observed over several years [Ashton et al., 2017]. More indirect impacts of rising temperatures and decreasing sea ice cover are shifts in the highly seasonal growth patterns of benthic communities [Morley et al., 2019, Barnes, 2017] or potentially changing predation regimes, i.e. due to the invasion of new species [Aronson et al., 2009]. Investigating such complex and uncertain interacting impacts in field studies is very difficult, especially in a remote location like the Antarctic shelf [Barnes et al., 2025].

In this study, we use a simulation model to investigate how the succession after an ice-scour is influenced by potential climate change effects, focussing on Bryozoan assemblages. Bryozoans are sessile, suspension-feeding organisms that grow in colonies on the seabed. They are a very common and species-rich taxon, which is particularly well represented on the Antarctic shelf [Barnes et al., 2014]. Additionally, bryozoan colonies compete for space by overgrowing each other, and these patterns of overgrowth are preserved in their shells. This offers the unique possibility of directly observing and quantifying competitive interactions from snapshots [Barnes and Rothery, 1996], which has lead to important insights into bryozoan community dynamics and successions: In general, polar bryozoan assemblages are very hierarchical [Barnes, 2002] and characterised by a strong trade-off between growth rate and competitive strength: strong competitors typically grow very slowly and establish new colonies at a low rate, but are able to overgrow and hence eventually exclude weaker competitors [Barnes et al., 2014]. Those weaker competitors, on the other hand, are able to compensate for competition losses by relatively fast growth and the ability to easily and rapidly colonise newly available space (disturbed areas) [Barnes and Clarke, 1998, Brown et al., 2004].

As is typical for Antarctic species [Arntz et al., 1994], Bryozoans show a clear seasonality in their dynamics. During winter, little light penetrates the cover of sea ice, which strongly limits primary productivity and thus food availability for benthic species [Peck et al., 2006, Robinson et al., 2021, Zwerschke et al., 2021]. As sea-ice cover decreases, primary productivity increases earlier, leading to shorter winters and longer growing seasons [Barnes, 2017, Morley et al., 2019]. Furthermore, weaker competitors become active relatively early during the growing season, while strong competitors often recruit later [Bowden, 2005]. Lengthening of the growing season can thus affect the successional dynamics between pioneer and climax species [Barnes et al., 2025]. Another factor modulating this response is the timing of icebergs: as it gets warmer and ice melts earlier, ice-scouring will likely occur earlier in the growing season. Finally, there might be further indirect effects on bryozoan assemblage dynamics due to changes in predation. Currently, common predators of bryozoans are echinoids (e.g. *Sterechinus neumayeri*), chitons (Polyplacophora), sea spiders (Pycnogona) and nudibranch gastropods. Grazing by these predators kills individual zooids and thus opens up empty space so that they can be seen as biological agents of physical disturbance [Barnes and Conlan, 2007]. Rising temperatures and sea ice loss affect all trophic levels [McClintock et al., 2008, Montes-Hugo et al., 2009, Trivelpiece et al., 2011], but making precise predictions about changes in predation intensity and type (e.g. through invading new predators, see [Aronson et al., 2009]) is currently not possible.

Our simulations describe how a bryozoan assemblage recovers after an ice-scour event. Each simulation starts with an empty surface of approximately 10 × 10 cm^2^, representing a small part of the seabed directly after the ice-scour hit. Individuals represent zooids of different species, which form colonies on the seabed and, on meeting each other, compete for space. Species-specific growth or colonisation rates are very difficult to measure and hence not available. Despite this scarcity of data, we aimed to capture the most characteristic properties of real-world assemblages: Species with high competitive strength are able to overgrow weaker species, but this competitive advantage comes at the cost of having relatively low colonisation rates, slow colony growth and a later onset of activity during the growing season. Mortality due to predation is modelled as the removal of zooids within small areas of the model world.

We conducted several simulation experiments, in which we contrasted a short growing season, representing current climatic conditions, with simulations in which the growing season has elongated due to rising temperatures. Additionally, we varied the timing of ice-scour events within the growing season, comparing icebergs that hit later (current state), with icebergs that hit earlier as temperatures rise. Finally, we included two predation scenarios, that were equal in intensity but differed in the spatial distribution of predation events, representing e.g. different types of dominant predators. We did not aim to make precise predictions, but to understand how the combination of these three factors shape successional dynamics and the coexistence of pioneer and climax species.

We hypothesised that an earlier timing of ice-scour during the growing season will disadvantage strong competitors during the early stages of assemblage development, as weaker competitors can benefit from their temporal advantage of having an earlier onset of activity (H1). We furthermore hypothesised that a longer growing season will change assembly composition by accelerating succession and hence shortening the period with high diversity, as strong competitors have more time to exclude weaker species (H2). Finally, we hypothesised that the spatial distribution of predation events further modulates assemblage dynamics. Predation events that are clustered in space instead of widely distributed could hit strong competitors disproportionately hard, as they have fewer colonies to regrow from and lower chances to recolonise disturbed areas (H3).

## 2 Methods

This study was conducted using a simulation model implemented in Netlogo (v6.4.0, [Wilensky, 1999]). All analyses were performed using R Statistical Software (v4.3.3) [R Core Team, 2023]. The code for reproducing the results is available as online supplementary material. In the following, we first provide a complete model description, following the Overview, Design concepts, Details (ODD) protocol for describing individual and agent-based models [Grimm et al., 2006, Grimm et al., 2010], before explaining how the model was analysed.

### 2.1 ODD Protocol

#### 2.1.1 Purpose

The model simulates the development of polar bryozoan assemblages shaped by seasonal growth, competition for space and external disturbances due to ice scouring and predation events. Sexually produced larvae can disperse through the water column, settle and metamorphose on rocky substrates. After successful recruitment, the initial zooid, termed an ancestrula, can colonise space by growing new zooids clonally. Colonies compete for space by attempting to overgrow each other (see Fig. 1 b and c). A competition-colonisation and a competition-growth trade-off is assumed, meaning that strong competitors (climax species) are able to overgrow weaker competitors (pioneers) at the cost of having lower colonisation and growth rates. Furthermore, the model incorporates seasonal species-specific growth, where strong competitors become active later in the growing season compared to good colonisers.

**Figure 1:**
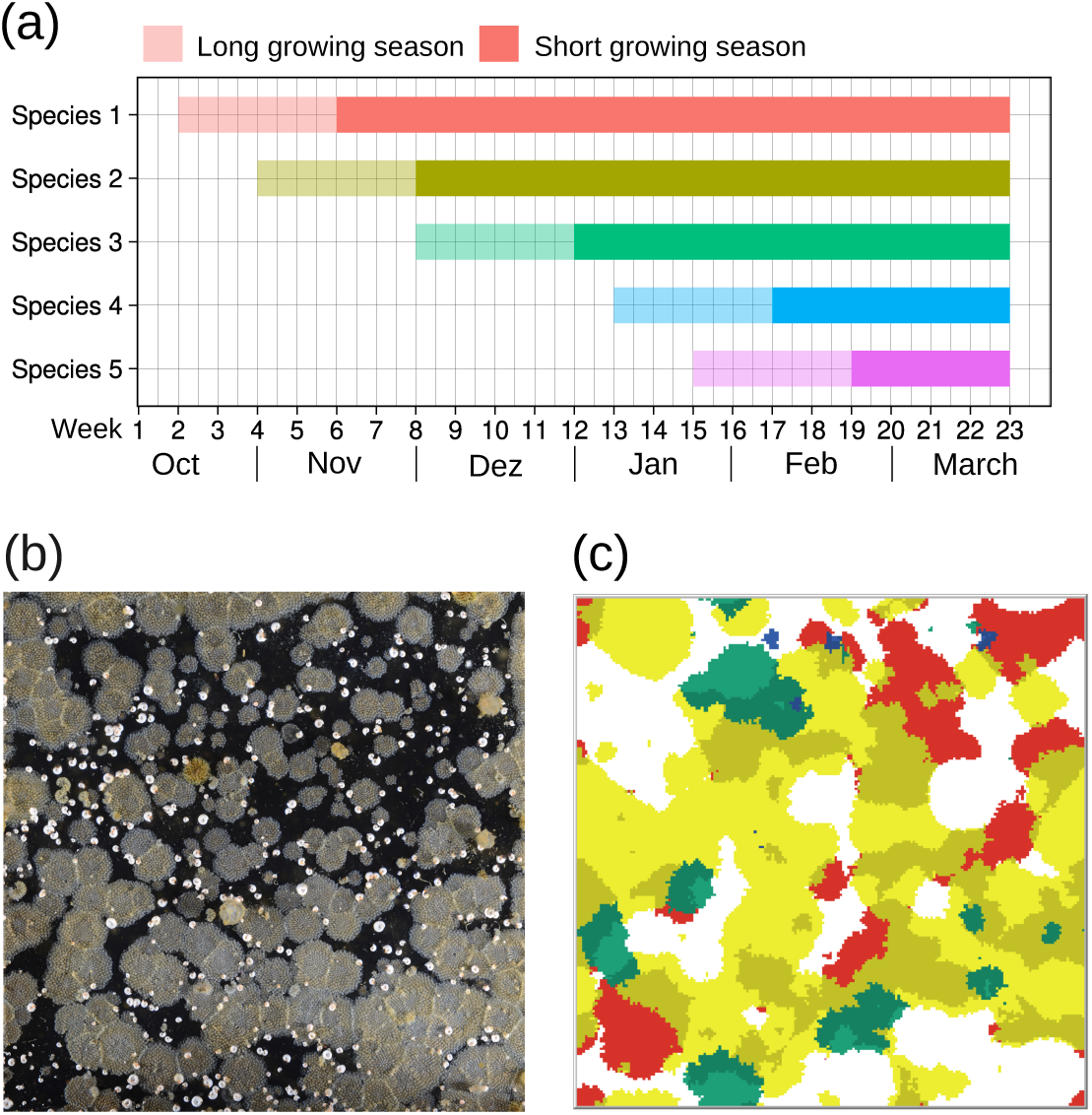
A simulation model of bryozoan assemblage dynamics. (a): Species-specific onset of activity in short and long growing season scenarios. We assume that species with higher competition strengths generally become active later during the growing season. For example, species 1 with competitive strength *c*_1_ = 0.1 is able to start colonising and growing after week 6 (mid November) in the short growing season scenario and after week 2 (mid October) in the long growing season scenario. By contrast, species 2 with competitive strength *c*_2_ = 0.2 is able to start after week 8 or 4, respectively. (b) Image of a real bryozoan assemblage (photo by: British Antarctic Survey, Cambridge, UK) (c) Screenshot of the model world depicting bryozoan colonies competing for space in a simulation with long growing season after 70 time steps. The simulation started in October, and the predation regime with few large disturbances was chosen. All other model parameters were chosen as in Tab. 2. Species 1=red, 2=yellow, 3=green, 4=blue and 5=purple (not yet established). Overgrown zooids have a slightly darker shade of the overgrowing species’ colour.

The model can be used to investigate how different disturbance regimes affect assemblage dynamics. Specifically, we look at a set of disturbance scenarios by varying three factors in the model: the timing of ice-scour events, the duration of the growing season as well as the spatial distribution of predation events. Insights from this model can be used to predict future biodiversity dynamics of benthic bryozoan assemblages under climate change.

#### 2.1.2 Entities, State Variables, Scales

The model world consists of a grid of 201×201 patches, wrapped in both dimensions. Each patch corresponds to approximately 0.5 x 0.5 mm of the surface of a submerged rock on the polar seabed (roughly corresponding to the size of an individual bryozoan zooid). This leads to a modelled area of 10×10 cm^2^. The size of the model world is based on the heated panels used in [Ashton et al., 2017]. Each grid cell can be occupied by a single living zooid, and each zooid has two state variables, namely *species ID* and *colony ID*. The former determines the species-specific competitive strength, colonisation rate and colony growth rate, as explained further below in the context of model initialisation. The latter is used to count abundances and to identify zooids on the edge of colonies, which is necessary to model colony competition. Each model time step represents the time a zooid needs to clone itself, which is roughly a week in polar environments.

#### 2.1.3 Process Overview and Scheduling

In total, the model includes four processes, which are executed at each time step and in the order indicated below. Note that the processes ***growth*** and ***colonisation*** can be executed in two different versions to capture a relatively short growing season, as currently observed, versus an extended growing season, as expected due to climate warming [Barnes et al., 2024b].

- The ***update-time*** procedure keeps track of the weeks, months and years that have passed since simulation start.
- The ***growth*** procedure describes how colonies grow in space. Zooids on the edges of colonies have the chance of spawning a clone on a neighbouring patch to increase the colony size. If the neighbouring patch is already occupied, the attacking zooid attempts to overgrow. All growth stops during Austral winter and restarts at species-specific time points in spring or summer.
- The ***colonisation*** procedure describes how new colonies are established on randomly selected patches. The colonising species is chosen based on its colonisation rate. Again, if the randomly chosen patch for colonisation is already occupied, the colonising zooid attempts to overgrow. Colonisation stops during winter and restarts at species-specific time points in spring or summer.
- The ***predation*** procedure simulates zooid mortality due to predation. The frequency and severity of these predation events can be varied.

#### 2.1.4 Design Concepts

The **basic principles** used in the model are (1) interference competition for a limiting resource, in this case space, (2) the concept of competition-colonisation and competition growth trade-offs and (3) competitive exclusion. Community structure, the abundances and spatial distribution of pioneer and climax species **emerges** from species **interactions**, in this case overgrowth competition between bryozoan colonies. Many processes have **stochastic elements**. For example, the location of colonisation and disturbance events are chosen randomly, as well as the neighbouring patch that is chosen during colony growth. Stochasticity also plays a role in the outcome of overgrowth competition. Note that colonies can be seen as an intermediate level of organisation, given that they consist of many zooids with the same *colony ID*, meaning that in principle they can be seen as **collectives**. However, all processes are implemented at the zooid level, so there is no colony-specific behaviour. Colony IDs are only relevant for **observation**, as they are needed to record abundances.

The design concepts **Adaptation**, **Objectives**, **Learning**, **Prediction** and **Sensing** do not play a role in this model.

#### 2.1.5 Initialisation

Each simulation run starts with an empty model world, which represents the state of the seabed right after an ice-scour. We do not simulate the ice-scour itself, which represents a mass mortality event in which all the colonies are scraped off the substrate. Thus, the state variables of all zooids (namely *speciesID* and *colonyID*) are initialised with the default value zero. The temporal variables *week* and *month* are initialised based on the chosen starting month. Selecting October as the starting month corresponds to an ice-scour in the last week of September. As we define October as the start of the year, this would result in the variables *week* and *month* both being initialised to 0. The last possible starting month is May, which corresponds to an ice-scour in the last week of April. Table 1 gives an overview over all model variables and their values at initialisation.

**Table 1:** A summary of all patch-specific and global variables used in the individual-based model.

| **Patch-specific model variables** |  |  |
| --- | --- | --- |
| Name | Description | Value at initialisation |
| species-ID | determines which species the zooid belongs to, used to determine competitive strengths as well as colonisation and growth rates | 0 (indicating an empty cell) |
| colony-ID | determines which colony the zooid belongs to, used to detect colony edges and abundances | 0 (indicating an empty cell) |
| **Global variables** |  |  |
| Name | Description | Value at initialisation |
| year | counts the number of years since the last ice-scour | 0 |
| week | determines current week within the year (0 = beginning of October; 47 = end of September) | 0,1,2...28 (depending on chosen starting month) |
| month | determines current month within the year (0 = October, 11 = September) | 0,1,2...7 (depending on chosen starting month) |

The model parameter *number-of-species* controls the size of the species pool. Each species in the species pool is assigned a competitive strength *c_i_* ∈ [0,1). The model parameter *competition-list* contains a list of all competitive strengths (see Table 2 for a list of all model parameters). For the study presented here, we chose a fixed pool of 5 species with the following fixed competitive strengths: 0.1 (species 1), 0.2 (species 2), 0.5 (species 3), 0.8 (species 4) and 0.9 (species 5). We chose those values because we wanted to cover a wide range of competitive strengths, as it is known that arctic bryozoan assemblages are very hierarchical with several pioneer species (here 1 and 2) and much stronger climax species (here: 4 and 5) [Barnes, 2002]. The species-specific colonisation rate *x_i_* and growth rate *r_i_* are then calculated via *x_i_* = 1 − *c_i_* and *r_i_* = 0.75 − 0.5*c_i_*, respectively.

**Table 2:** A summary of all parameters used in the individual-based model and the values used for our simulation experiments.

| Parameter name | Description | Standard values for this study |
| --- | --- | --- |
| number-of-species | size of the species pool | 5 |
| competition-list | contains competitive strength of each species in the species pool, chosen by us | [0.1, 0.2, 0.5, 0.8, 0.9] |
| colonisation-list | contains colonisation rate of each species in the species pool, calculated based on competitive strengths | [0.9, 0.8, 0.5, 0.2, 0.1] |
| growth-list | contains growth rate of each species in the species pool, calculated based on competitive strengths | [0.7 0.65 0.5 0.35 0.3] |
| overgrowth-threshold | difference in competitive strengths that leads to overgrowth (instead of a tie) | 0.1 |
| overgrowth-stochasticity | boundary of the distribution used to add stochasticity to competitive outcomes | 0.2 |
| max-colonisation-events | Number of colonisation attempts per time step | 5 |
| predation-number | number of circular areas cleared by predation | 1 or 9 |
| predation-radius | radius of each circular area cleared by predation | 15 or 5 |

High competitive strengths thus translate into low colonisation and growth rates, and vice versa. Our reasoning for choosing these values is twofold: Based on empirical observations, we expect that the factor between fastest and slowest growth rate in the system to be roughly 3 and that differences in colonisation rates are much higher compared to differences in growth rates.

#### 2.1.6 Input Data

There is no input data.

#### 2.1.7 Submodels

##### Update-time

During this procedure, temporal variables are updated to keep track of how the number of model time steps translates into weeks, months and years. As each time step is considered to represent one week, a month consists of four time steps and a year of 12 months or 48 weeks.

##### Growth

This submodel describes how colonies grow in space through cloning of individual zooids located on the edge of the colony. Zooids can only clone themselves when they have enough resources, which is not the case during polar winter. Additionally, we assume that a species’ onset of activity in spring is related to their competitive strength, with stronger competitors requiring more resources and thus becoming active later. At each time step, all zooids that can potentially produce a clone are identified. These zooids must (1) belong to a species that is active in a given week (see Fig. 1 (a) for an overview of when species become active) and (2) be located on the edge of their colony.

Zooids located on the edge of a colony differ from at least one of their four direct neighbours in the value of the state variable *colony ID*, meaning they are either occupied by a different colony, or empty. For each of the edge zooids, we then select one of these neighbours at random. With a probability that is given via the species-specific growth rate *r_i_*, we assume that the edge zooid attempts to establish a clone on the selected neighbouring patch. Two cases can occur: First, if the target patch is empty, then we assume that the attempt is automatically successful, meaning that the state variables *colony ID* and *species ID* of the target patch are updated with the values of the corresponding state variables of the edge zooid. Second, if the target patch is already occupied by another zooid, we assume that the selected zooid attempts to overgrow.

The success of an overgrowth attempt is determined based on the ratio of both species’ competition strengths, *c_i_/c_j_*, which is larger than 1 if the attacking zooid (*i*) is a stronger competitor compared to the defender (*j*), and smaller than 1 in the opposite case. We consider the overgrowth attempt successful, if the following condition is fulfilled:

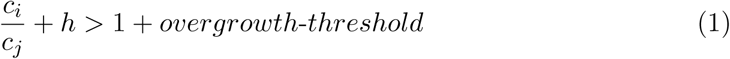

This is based on empirical observations on competitive outcomes: In general, competition is very hierarchical, and the stronger competitor wins the large majority of inter-actions [Barnes, 2002]. However, weaker zooids can occasionally also overgrow stronger ones [Barnes, 2002]. To enable the presence of these competitive reversals, we add some stochasticity to the competitive outcomes by adding a random number *h* between 0 and *overgrowth-stochasticity*. Additionally, interactions between species with very similar competitive strengths (which includes most intraspecific interactions) often have undecided outcomes, which means that both colonies cease growth in that direction and a static boundary forms [Tanaka and Nandakumar, 1994, Barnes and Rothery, 1996]. The parameter *overgrowth-threshold*determines the difference in competitive strength that is necessary to produce a clear overgrowth. Currently, we lack sufficient data to be able to choose these parameters based on observations. However, we found that varying the parameter values did not affect the outcome of our simulations (see Supplementary Figure 4).

If the overgrowth attempt is successful, then the state variables *colony ID* and *species ID* of the target patch are updated with the values of the corresponding state variables of the attacking zooid.

##### Colonisation

This procedure describes how new colonies are established. We do not explicitly model the production and dispersal of larvae, assuming that there are other assemblages outside our model world that serve as a constant source of larvae. As with growth, species only have enough resources to produce larvae and hence establish new colonies during their species-specific growing season [Barnes and Conlan, 2007, Bowden, 2005].

To model the establishment of new colonies, we first randomly select *max-colonisation events* patches in the model world (see Supplementary Fig. 3 for the effect of using different values of max-colonisation events). For each of these patches, one of the species that is currently active (see Fig. 1 a) is chosen with a probability that reflects that species’ colonisation rate. Species with a high colonisation rate get chosen more often and are hence quicker at establishing new colonies. The chosen species then attempts to colonise the empty patch. Three cases occur: First, if the target patch is empty, colonisation is always successful. Second, if it is already occupied by a different colony of the same species, colonisation fails. Third, if it is occupied by a colony from a different species, the colonising zooid attempts to overgrow the existing zooid, exactly as explained above in the context of the growth procedure. After a successful colonisation or overgrowth attempt, a new colony is established and both state variables of the target patch are updated accordingly.

##### Predation

This procedure mimics zooid mortality due to predation, e.g. by sea urchins. The frequency of predation events is controlled by the model parameter *predation-number*, while their intensity in terms of affected area is controlled by the parameter *predation-radius*. At each time step, we first randomly select *predation-number* patches and then remove all zooids inhabiting patches in a distance smaller than *predation-radius* to that target patch by setting their state variables *species-ID* and *colony-ID* to zero.

### 2.2 Disturbance scenarios and analysis

We used the simulation model described above to simulate a range of scenarios in which we varied (1) the length of the growing season, (2) the timing of ice-scour events and (3) the spatial distribution of predation events.

In particular, we first contrasted simulations with a short growing season (November-April), representing the current climate, and those with an extended growing season (October-May), which we expect as a result of climate change [Barnes, 2017]. The timing of ice-scour was then varied by changing the starting-month of the model run. In general, we expect that as temperatures rise, icebergs will hit earlier in the growing season. In the case of a short growing season, we considered 6 possible starting months (namely November, December, January, February, March or April). By contrast, in the extended growing season, we additionally considered October and May as possible starting months.

We furthermore considered the impact of potential changes in predation. While it is known that climate change will affect all trophic levels in the Southern Ocean [McClintock et al., 2008, Trivelpiece et al., 2011], precise predictions of how predation on bryozoans will change are so far not available. Still, predation opens up additional empty space for colonisation and might thus strongly affect the course of succession in bryozoan assemblages. It can be expected that simply changing the frequency or intensity of predation will affect successional patterns in predictable ways, in line with the intermediate disturbance hypothesis. This was confirmed in a preliminary analysis (see Supplementary Figure 2), which also revealed that the spatial resolution of these predation events is important. In the following, we therefore focused on two specific scenarios. In the first scenario, we simulated few large disturbance events with only one predation event per week that cleared a relatively large space (*predation-number* = 1 and *predation-radius* = 15, as in Fig. 1). In the second scenario, we included many small predation events (*predation-number* = 9, *predation-radius* = 5). In both scenarios, the total area cleared per time step was comparable (based on the area of a circle: 1 · *π* · 15^2^ = 9 · *π* · 5^2^ ≈ 707), even though the realised number of patches was occasionally slightly lower if areas of impact overlapped.

Combining all three factors resulted in a total number of 14 + 14 = 28 disturbance scenarios. Every scenario was replicated 10 times, resulting in a total number of 280 simulations. During each simulation, the number of zooids per species and the number of colonies per species were tracked over time. All simulations ran for 500 weeks, which is roughly ten years in the model world. All other parameters not varied in the scenarios were chosen as in Tab. 2.

## 3 Results

### 3.1 General pattern of succession

Regardless of the specific scenario (considering starting month, season length and predation regime, see section 2.2), assemblage development in our model was always characterised by the same step-wise pattern of succession, as shown in Fig. 2. Note that only two possible starting months are shown, to visualise the most important differences between scenarios that started at the beginning of the growing season (November) versus scenarios that started in the middle of the growing season (January). Scenarios that started late in the season are not discussed here, as they showed very similar patterns compared to scenarios starting early, due to the effective loss of the first growing period and the absence of activity during winter. Corresponding figures showing the number of colonies instead of the number of zooids can be found in Supplementary Figure 1.

**Figure 2:**
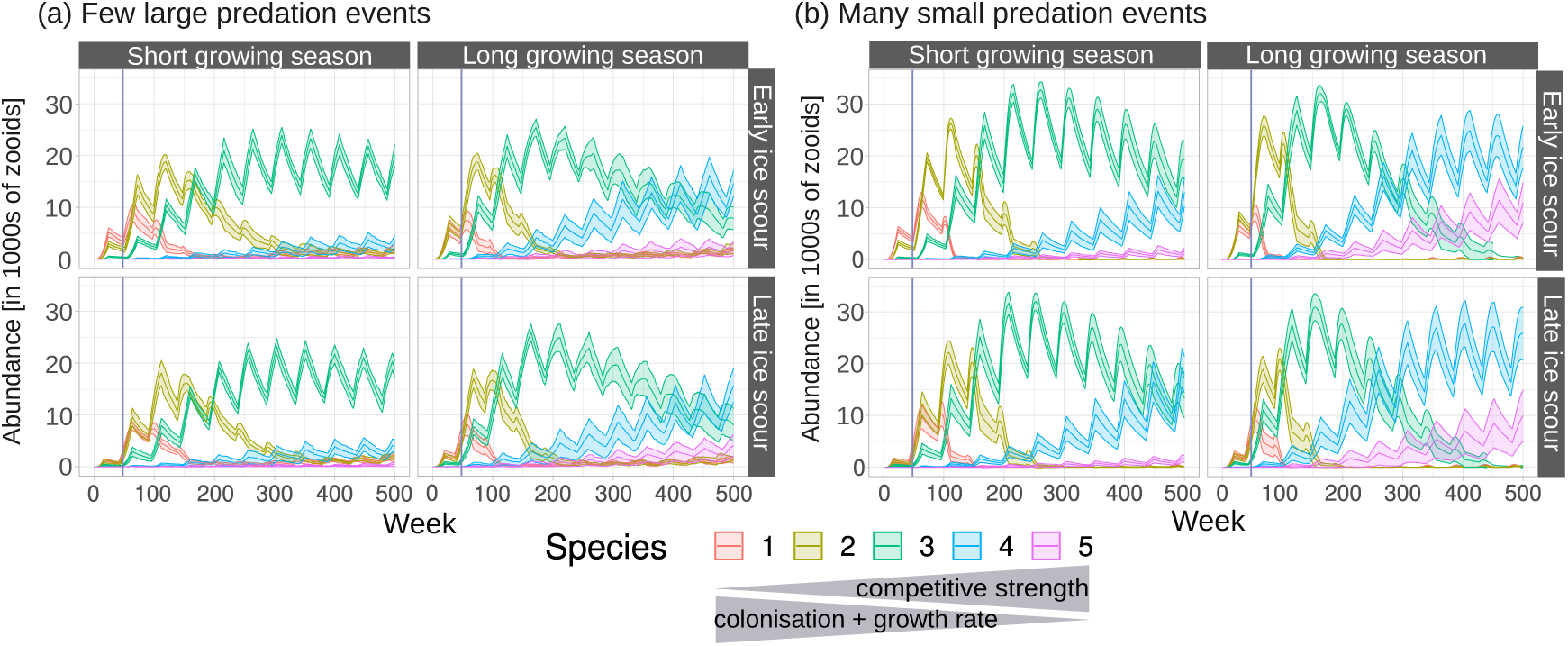
Successional dynamics in bryozoan assemblages under varying disturbance regimes. We simulated impacts of changes in disturbance regime by varying (1) the length of the growing season, (2) the timing of ice-scour events within the growing season and (3) the spatial distribution of predation events. Short growing season (left panels) refers to a current growing season from Nov-April, while long season (right panels) refers to an extended growing season due to climate warming (Oct-May). Upper panels show dynamics after an ice-scour hit early in the growing season (Starting Month: November) while lower panels show dynamics after an ice-scour hit in the middle of the growing season (Starting Month: January). Finally, we considered two scenarios of predation, one with few, large events (a) and a second one with many smaller predation events (b). All other parameters were chosen as in Tab. 2. Data shown is the mean of 10 replicates, shaded areas correspond to standard deviation. The vertical lines indicate abundances after exactly one year (48 weeks) as shown in Fig. 4

The first species that settled on the empty seabed at the beginning of the simulation is a pioneer species (species 1 in red, with lowest competition strength but highest colonisation and growth rates). Initially, the abundance of this species grew quickly due to the high availability of uninhabited space and hence the lack of competition. However, its abundance also declined quickly soon after reaching a maximum value, as the amount of available space decreased and competition intensified. Following this, species 2 (in yellow) was the most abundant one, then species 3 (in green), followed by species 4 (in blue) and finally species 5 (in purple). The described dynamics are superimposed by recurrent drops in abundances, which occurred during the winter months when all bryozoan activities stopped but predation continued.

The replacement of dominant species was relatively fast at the beginning of the simulation but slowed down during simulation runtime. This is due to the assumed competition-colonisation and competition-growth trade-offs: each species was replaced by the next best competitor because of that species’ competitive advantage, which comes at the cost of having slower colonisation and colony growth rates, so that the whole dynamics decelerated over time. The replacement could eventually become so slow, that the dominance of species 4 and 5 was not observed within a simulation runtime of 500 weeks. We never theless decided against even longer simulation runtimes because empirical data suggests that it is unlikely for a seabed area to remain undisturbed by an ice-scour for more than five years [Deregibus et al., 2017, Barnes et al., 2024b].

### 3.2 Speed of succession depends on season length and predation regime

We found that longer growing seasons lead to an accelerated succession (compare left and right panels in Fig. 2 a and b, respectively). This result was expected, given that longer growing seasons directly translate into more active weeks per year, that is more time for colonisation, colony growth and competition between colonies. A short season length implies that the number of weeks in which all species are active is low, so that pioneer species have relatively many weeks in which they hardly experience spatial competition. The longer the season length, the more biomass loss due to competition is expected because those species that act as strong competitors have more time to colonise and exclude others.

An accelerated succession also occurred with smaller but more frequent predation events (compare panels a and b in Fig. 2), which was coupled with overall higher abundances. This effect of the predation scenarios was at first sight surprising, given that the number of patches that have been cleared per week is almost identical (see section 2.2). The observed effect must therefore arise from the spatial distribution of predation events. Larger areas take longer to be refilled by colony growth from the side, meaning there is more time for recolonisation, which mostly benefits pioneer species. They can easily establish new colonies on the cleared ground without experiencing much competition. Small and widely distributed areas, on the other hand, are not only quickly filled by colony growth from the side, but they are also less likely to kill whole colonies, which mostly benefits climax species.

We furthermore found that the effects of season length and predation regime superimposed, meaning that the fastest succession was observed with many small predation events in combination with a long growing season (right panels in Fig. 2 b). The slowest succession, on the other hand, occurred with few but large predation events in combination with a short growing season (left panels in Fig. 2 a). By contrast, the exact timing of an ice-scour and hence the starting month of the simulation had only a minor impact on the speed of succession (compare top and bottom panels in Fig. 2 a and b, respectively).

Note that, as a result of these impacts on successional speed, pioneer and climax species benefitted from different scenarios. Pioneer species 1 and 2 survived longest in scenarios with a short season length in combination with few but large predation events (Fig. 3 a and b). Climax species 4 and 5, on the other hand, benefitted in scenarios with a long season length in combination with many small predation events (Fig. 3 c and d).

**Figure 3:**
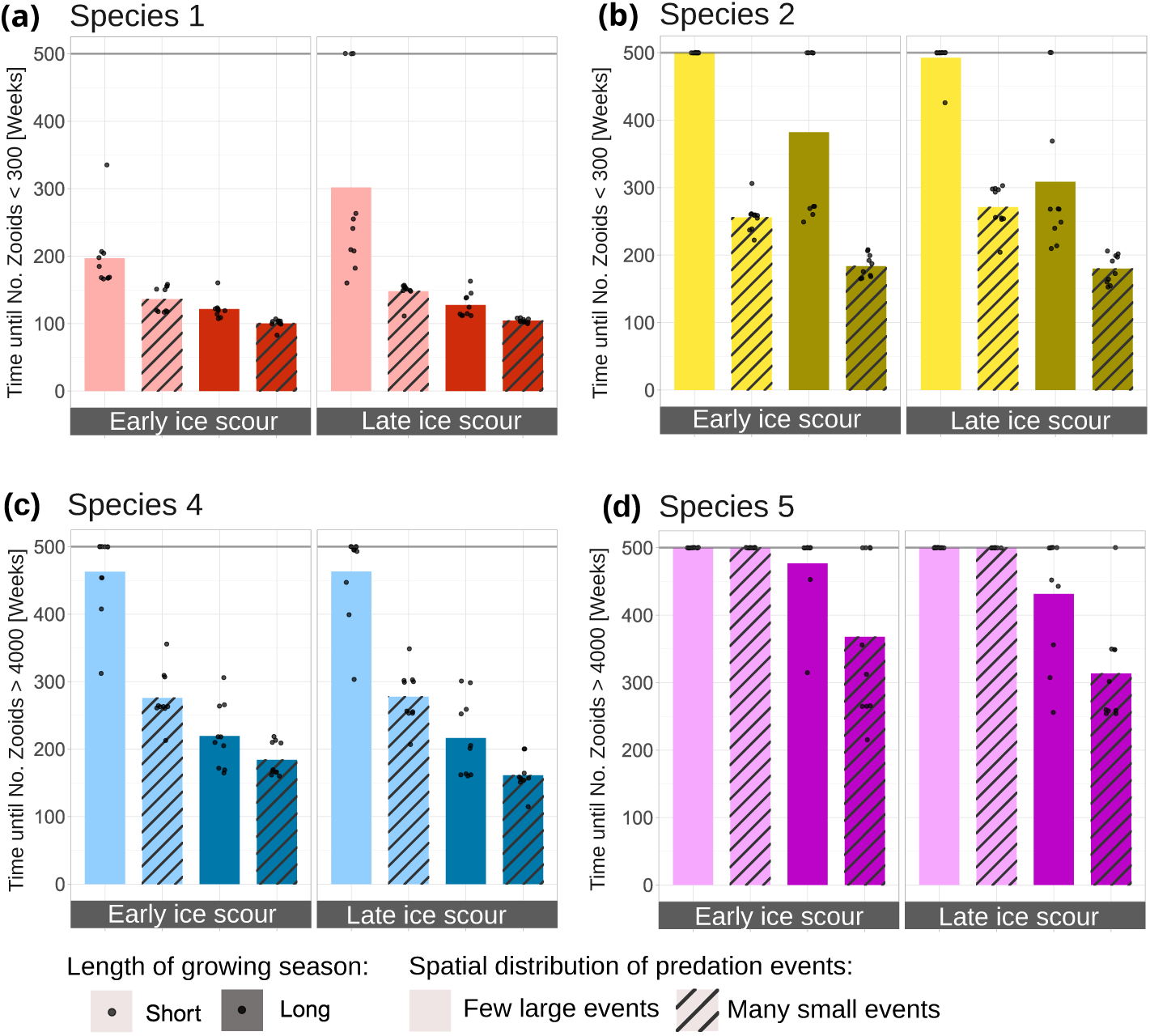
Key time points in the dynamics of pioneer and climax species under varying disturbance regimes. For pioneer species 1 (a) and 2 (b) we compared the number of time steps until they were almost excluded by stronger competitors. Due to ongoing (re-)colonisation, we never actually observed extinction. Instead, we considered a threshold of 300 zooids as the time point where a species becomes irrelevant for overall assemblage dynamics. For climax species 4 and 5, we considered the time points at which they first surpass the threshold of 4000 zooids. Predation scenarios are as in Fig. 2. For both cases, if the threshold was not reached within the simulation runtime of 500 weeks, the plot shows 500 as the key time point. Data points represent individual simulation runs, bar plots represent the mean values.

### 3.3 Timing of ice-scour events affects early assemblage development

Although the overall pattern and speed of succession was almost unaffected by the timing of the ice-scour, it did affect short-term dynamics at the beginning of the simulation. This becomes apparent when looking at the mean abundances of all species after one year, as shown in Fig. 4. In general, earlier starting months showed lower abundances, as these correspond to assemblages that just went through winter when many zooids were lost due to predation. Total abundances thus increase with later starting months, and this increase is stronger in the longer growing season (right panels).

**Figure 4:**
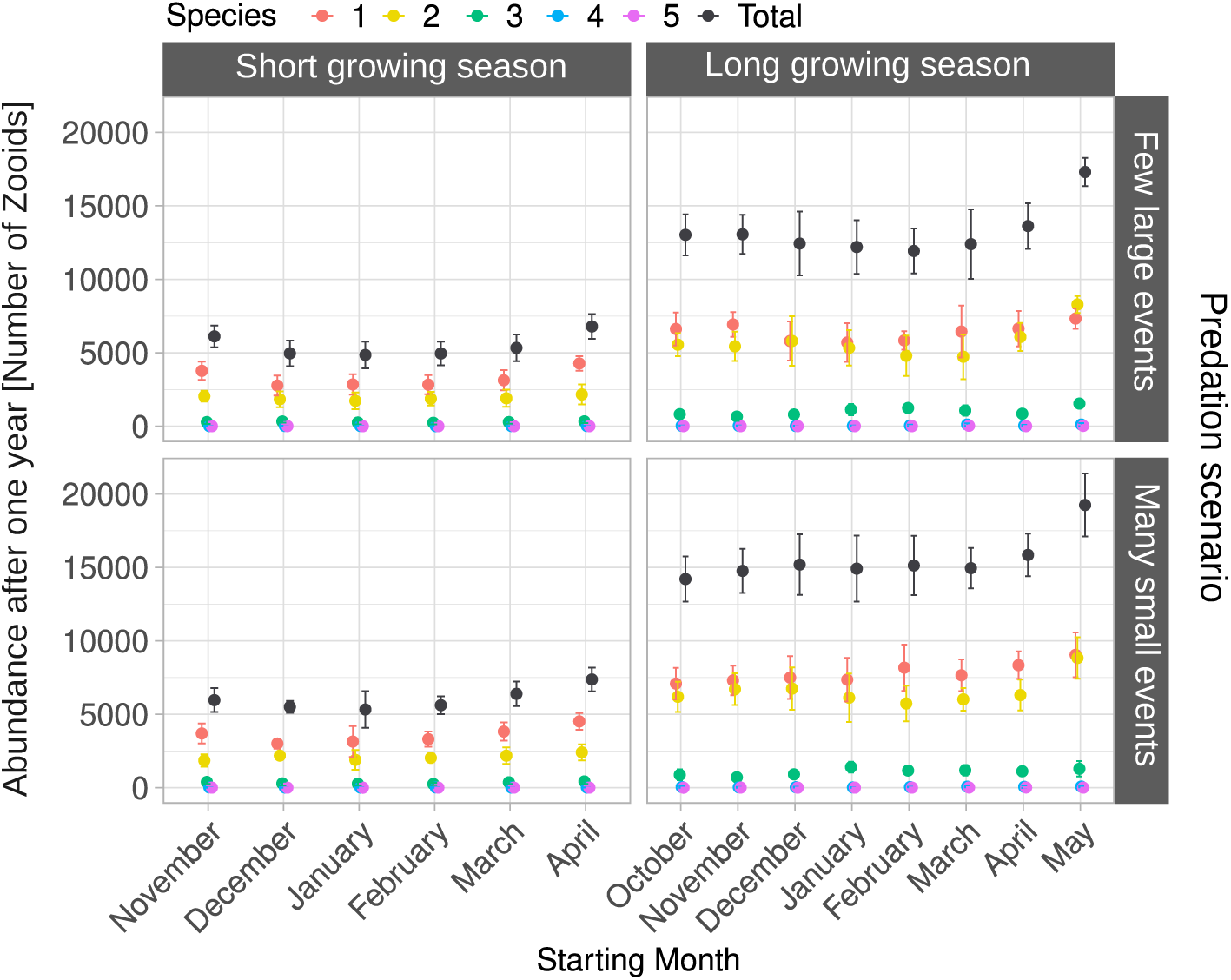
Timing of ice-scour events affects abundances in the early stages of assemblage developments. Abundances are shown for 1 year after the simulation start and thus the ice-scour event. The effect of timing of icescouring is compared in simulations with a short season length (left panels) and an extended season length due to climate change (right panels), as well as for our two different predation regimes (upper vs. lower panels).

In short season simulations, we additionally found that when ice-scours hit in the middle of the growing season, this had a small negative effect on the abundance of the pioneer species 1. This was because weaker competitors lose their temporal advantage of having an earlier onset of activity when icebergs hit in the middle of the season. Overall, we thus saw a U-shaped relation between abundance and starting month in simulations with a short growing season. The negative effect of starting in the middle of the growing season became less strong when season length is increased and when many small predation events are considered.

## 4 Discussion

The goal of this study was to investigate how successional dynamics in bryozoan assemblages are affected by complex changes in disturbance regimes due to climate-driven sea ice loss. Using a model of bryozoan assemblage dynamics, we ran simulation experiments in which we varied the timing of ice-scouring, the length of the growing season and the spatial pattern of predation. Our simulation results suggest that the composition of a given bryozoan assemblage is not only tightly linked to its age since last disturbance by ice-scour, as expected [Brown et al., 2004, Barnes et al., 2014, Robinson et al., 2021], but also to the length of the growing season and to the spatial distribution of predation events.

We expected earlier ice-scouring to benefit pioneers during early successional stages (H1) and could confirm this in simulations with a short growing season, but not in those with a longer growing season. However, our results confirmed H2 and H3. Fast succession occurs in scenarios with many small predation events in combinations with a long growing season, while slow succession can be observed in scenarios with few large predation events and a short growing season. It is likely that we did not see pioneers benefitting from earlier ice-scours in scenarios with a longer growing season because the potential advantage from earlier onset of activity was not strong enough to compensate the negative effect of a longer growing season. In general, the effects of the timing of ice-scours were small and only appeared in very early stages of assemblage development.

Our simulations allow us to derive predictions about the effect of changing ice-scour frequencies, even though we did not explicitly vary this factor in our scenarios. In our model, frequent ice-scouring translates into short simulation runtimes and hence dominance of pioneer species. Rare ice-scouring, on the other hand, translates into long simulation runtimes that allow for competitive exclusion and hence dominance of climax species. Thus, we can expect that bryozoan richness (and diversity) on the shallows of the Antarctic shelf is highest in areas that have a moderate rate of ice-scouring, which is in line with previous empirical work [Barnes, 1999, Barnes and Conlan, 2007, Smale et al., 2008, Zwerschke et al., 2021, Robinson et al., 2021] and the intermediate disturbance hypothesis [Connell, 1978, Wilkinson, 1999, Roxburgh et al., 2004, Miller et al., 2011, Sheil and Burslem, 2013, Moi et al., 2020].

In order to keep the model structure and computing requirements manageable, despite the multi-level complexity of the empirical system, we had to make several simplifying assumptions. This is particularly true for constant colonisation rates, which reflect the assumption that recruitment happens from sites that are relatively far away and hence not affected by changing conditions, or at least not simultaneously. A more realistic model version would be based on a much larger model world, resembling trophic meta-communities [Gross et al., 2020], in which colonies are able to produce larvae that can travel from one assemblage to another. Ice-scour disturbances would then affect only part of the model world, leading to a high heterogeneity of successional stages, as observed in the real world [Gutt, 2001]. Dispersal of benthic larvae was previously investigated in a modelling study by [Potthoff et al., 2006], who found that the minimum dispersal distance required for persistence was smaller than expected. This model could be used as a starting point to extend our own model to include more realistic dispersal mechanisms. However, building and analysing such an extended model version would go far beyond the scope of this study.

Due to its simplicity, and because parameters have not been calibrated based on empirical data, our model is not well suited to make quantitative predictions of future bryozoan assemblage dynamics. This is mostly due to a lack of empirical data for the observed competition-colonisation and competition-growth trade-off functions. Collection of such data in polar nearshore environments is expensive and logistically challenging, partly because it requires SCUBA to recover substrata with competing assemblages on. Very few polar research stations have SCUBA facilities and of those that do, only two also monitor ice-scouring (Carlini and Rothera, [Deregibus et al., 2017]). However, we ran several additional simulations in which we varied key parameter values (see Supplementary Figures 2-4), which showed that the general patterns observed in our study (i.e. how the factors we varied affected the speed of succession by favouring either pioneer or climax species) do not depend on exact parameter values.

Despite the discussed simplifications, we can derive qualitative predictions for future changes in observed large-scale biodiversity patterns based on our simulation results. Short-term climate-change scenarios involve an increased frequency of ice-scours [Barnes et al., 2014, Barnes et al., 2024b], so on average, bryozoan assemblages would be younger. In the shallows, this means that very diverse systems might turn into deserts dominated by very few pioneers [Barnes et al., 2014, Brown et al., 2004]. Such a rapid loss of diversity would also imply a loss of hierarchy in the system, which could further threaten those systems, given that competitive hierarchies in bryozoan assemblages have been shown to mitigate network instability by keeping destabilising feedback loops relatively weak [Koch et al., 2023, Koch et al., 2024]. Our simulation results suggest that this trend could be counteracted by an extended growing season, which benefits climax species and leads to overall increased abundances. Along similar lines, intermediate or long-term climate-change scenarios involve low frequencies or even the absence of ice-scours [Cook et al., 2005], which would again benefit climax species. Here, an extended growing season could have an additive or even amplifying effect. Taken together, this means that extended growing seasons might first mask the effect of more frequent ice-scours and then accelerate the effect of less frequent ones, so that observed changes in assemblage composition over time might occur more rapidly than previously expected.

Potential changes in predation patterns are another factor that can modulate successional dynamics in bryozoan assemblages. Our results suggest that, depending on the spatial distribution of predation events, either pioneer or climax species can be favoured by such changes. It is well-known that sea ice plays a critical role in structuring antarctic ecosystem dynamics [Trivelpiece et al., 2011] and that a loss of sea ice cover due to global warming has impact on all levels of the food web [McClintock et al., 2008, Morley et al., 2019]. These effects are usually studied with a focus on understanding the consequences for upper trophic levels (e.g. the effect of phytoplankton blooms on krill and penguin populations [Montes-Hugo et al., 2009, Trivelpiece et al., 2011]) while our understanding of impacts on lower trophic levels, including benthic communities, remains very limited. In general, it is assumed that higher water temperatures can promote the spread of non-indigenous cold-water species [Barnes and Conlan, 2007], and some studies have hinted at potential invasions of shell crushing predators such as crabs [Aronson et al., 2009]. In this study, we chose two very general predation scenarios in which we compared the spatial distribution of predation events, which could for example reflect a change in the dominant predator species. Our results indicate an effect of the predation scenario on successional speeds, suggesting that it is worthwhile to study changes in predation regimes in more detail, for example by extending the individual-based model to include other potential predator impacts.

Our study constitutes an important step towards understanding how benthic biodiversity will be impacted by climate change. Our simulations revealed that not only the frequency of ice-scours is important for the maintenance of benthic biodiversity but also other factors that modulate the biotic interactions in this disturbance-driven ecosystem, like the length of the growing season or the spatial distribution of predation events. This interplay between complex changes in physical disturbance regimes on the one hand and biotic interactions on the other is likely to also play a role in many other disturbed ecosystems. To understand combinations of different climate change impacts on higher levels of biological organisation like communities and ecosystems, such biotic interactions necessarily need to be considered [Orr et al., 2020], which highlights the need for an ecological network approach to multiple stressors research [Bruder et al., 2019]. Our study demonstrates that using simulation models that allow us to assess the impact of different types of climate change effects, both individually and in combination, can be valuable tools in this context.

## Acknowledgements

Previous model versions have been studied by T. H. M. Ritter and L. S. Krüger as part of their M.Sc. theses. We thank the whole department of Eco-Evolutionary Modelling and in particular Felix Jäger for insightful discussions and helpful feedback on the manuscript. KTA and FK acknowledge funding through the Deutsche Forschungsgemeinschaft (DFG, project number 451967415 (AL 2563/1-1)).

## Author contribution statement

MB: Conceptualization, Formal analysis, Investigation, Methodology, Software, Visualization, Writing – original draft, Writing – review and editing

KTA: Conceptualization, Funding acquisition, Investigation, Methodology, Project administration, Supervision, Writing – original draft, Writing – review and editing

DKAB: Conceptualization, Methodology, Validation, Writing – review and editing

FK: Conceptualization, Formal analysis, Investigation, Methodology, Project administration, Software, Supervision, Writing – original draft, Writing – review and editing

