## Supplementary Material for "Forecasting bryozoan assemblage dynamics under simulated climate change"

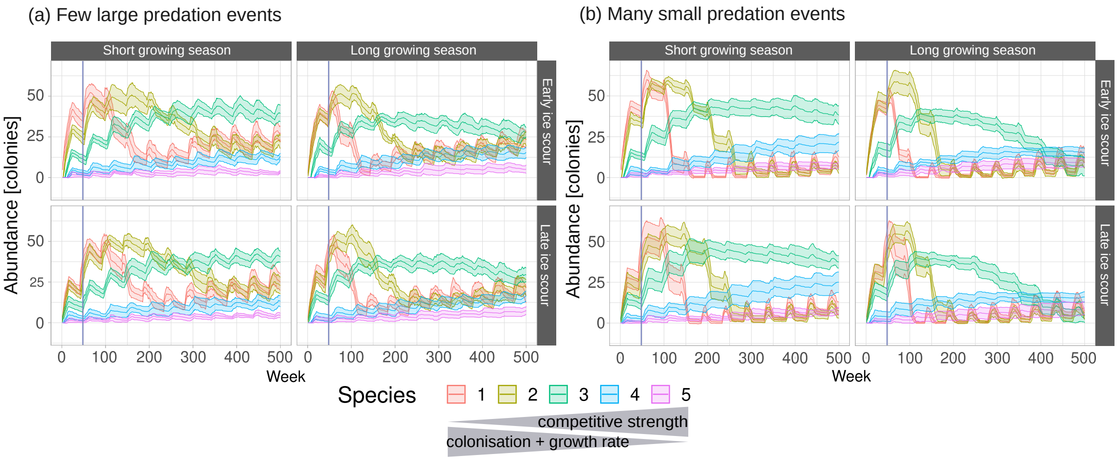


Supplementary Figure 1: **Successional dynamics in bryozoan assemblages under varying disturbance regimes.** Same simulation results as in Figure 2 in the main text, but abundances are given in number of colonies instead of number of zooids. Short growing season (left panels) refers to a current growing season from Nov-April, while long season (right panels) refers to an extended growing season due to climate warming (Oct-May). Upper panels show dynamics after an ice-scour hit early in the growing season (Starting Month: November) while lower panels show dynamics after an ice-scour hit in the middle of the growing season (Starting Month: January). Finally, we considered two scenarios of predation, one with few, large events (a) and a second one with many smaller predation events (b). All other parameters were chosen as in Tab. 2. Data shown is the mean of 10 replicates, shaded areas correspond to standard deviation. The vertical lines indicate abundances after exactly one year (48 weeks) as shown in Fig. 4 in the main text

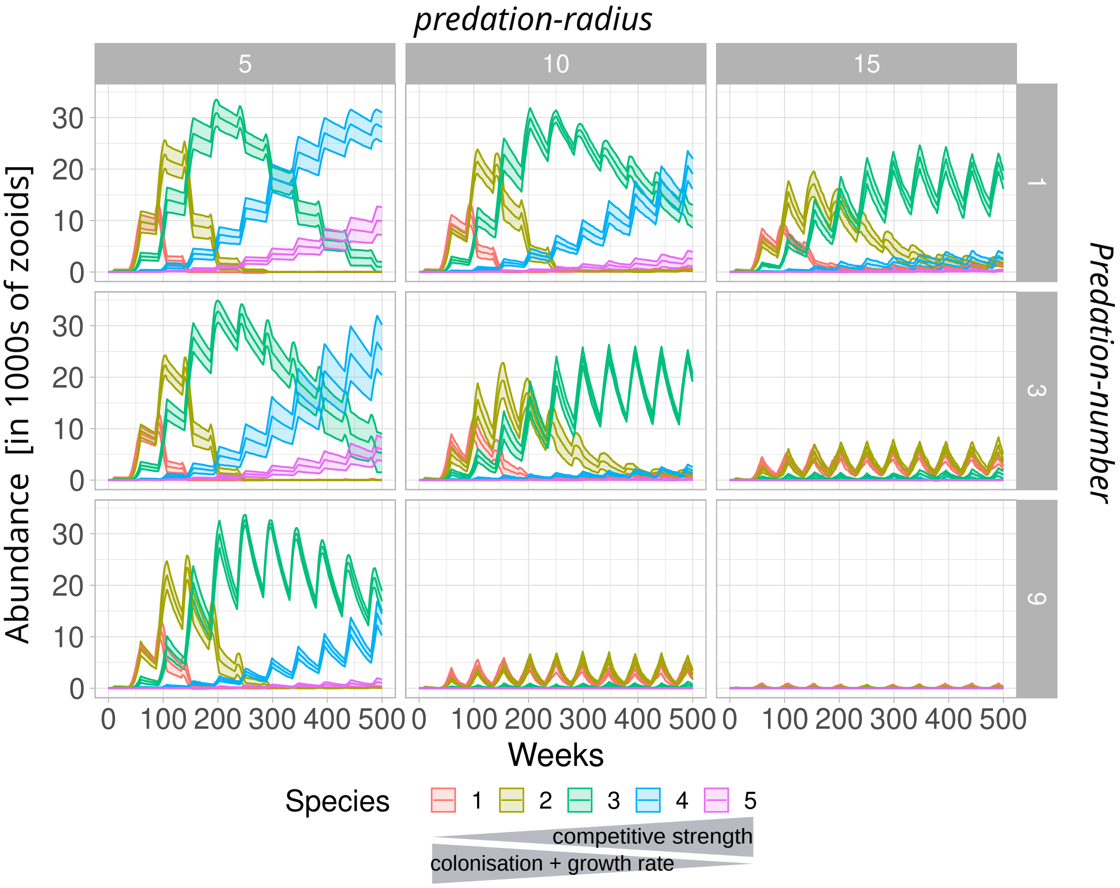


Supplementary Figure 2: **Successional dynamics with varying frequency and intensity of predation events.** Simulations were run with varying *predation-radius*, determining the size of area cleared by each predation event and *predation-number*, determining the number of predation events per time step. As predicted by the Intermediate Disturbance Hypothesis, higher intensity of disturbance through predation promotes pioneers and at some points prevents any species from being able to establish themselves. All simulations shown were run with a short season length and starting month was set to February. Data shown is the mean of 10 replicates, shaded areas correspond to standard deviation.

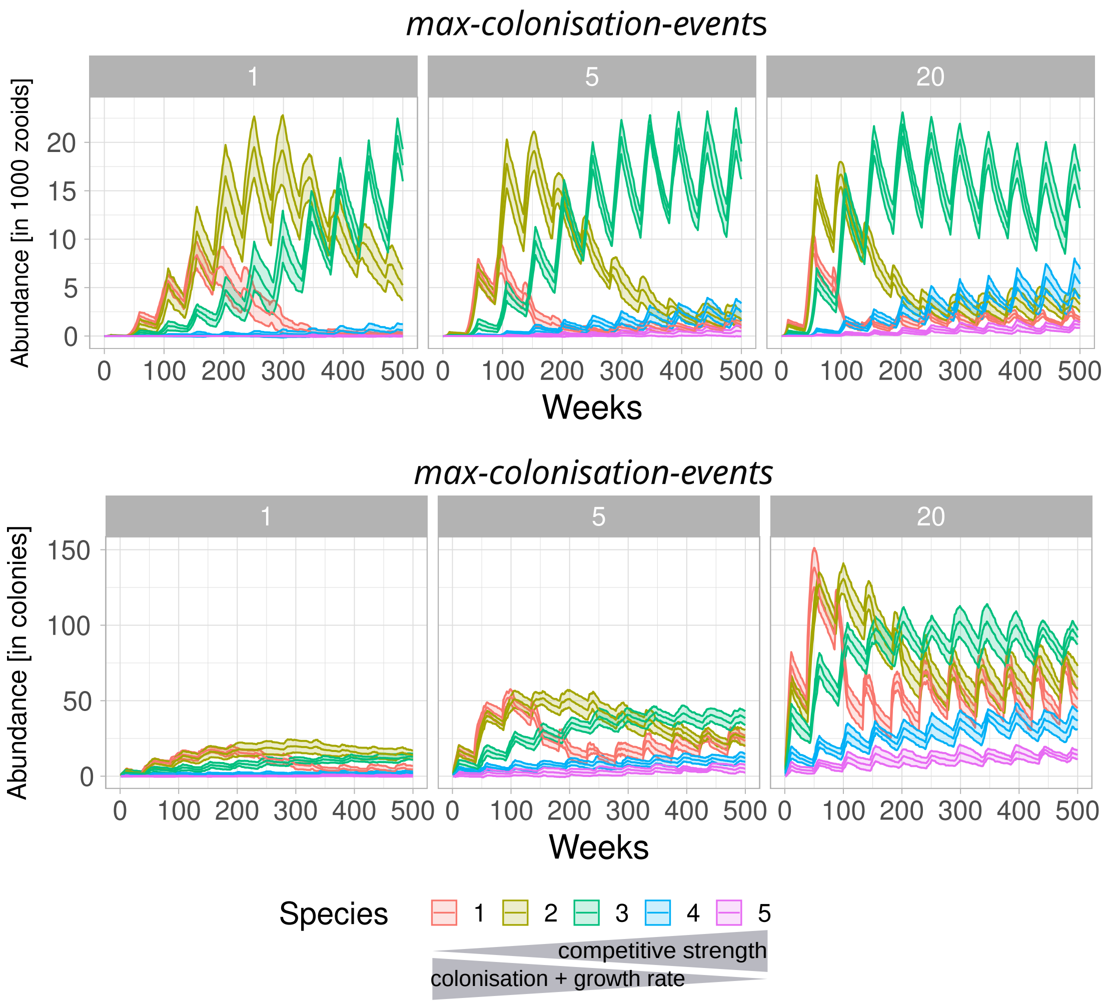


Supplementary Figure 3: **Successional dynamics with varying *max-colonisation-events*.** The parameter *max-colonisation-events* the maximum possible number of colonisation events that are possible per time step. The main analysis is based on *max-colonisation-events* = 5. Here, we show that varying this value has no effect on the qualitative results of our study. Higher values will speed up the successional dynamics, but they affect pioneers and climax species equally. We see a stronger effect when we count abundances in terms of colonies, as higher *max-colonisation-events* lead to more but smaller colonies. All simulations shown were run with a short season length and starting month was set to February. Data shown is the mean of 10 replicates, shaded areas correspond to standard deviation.

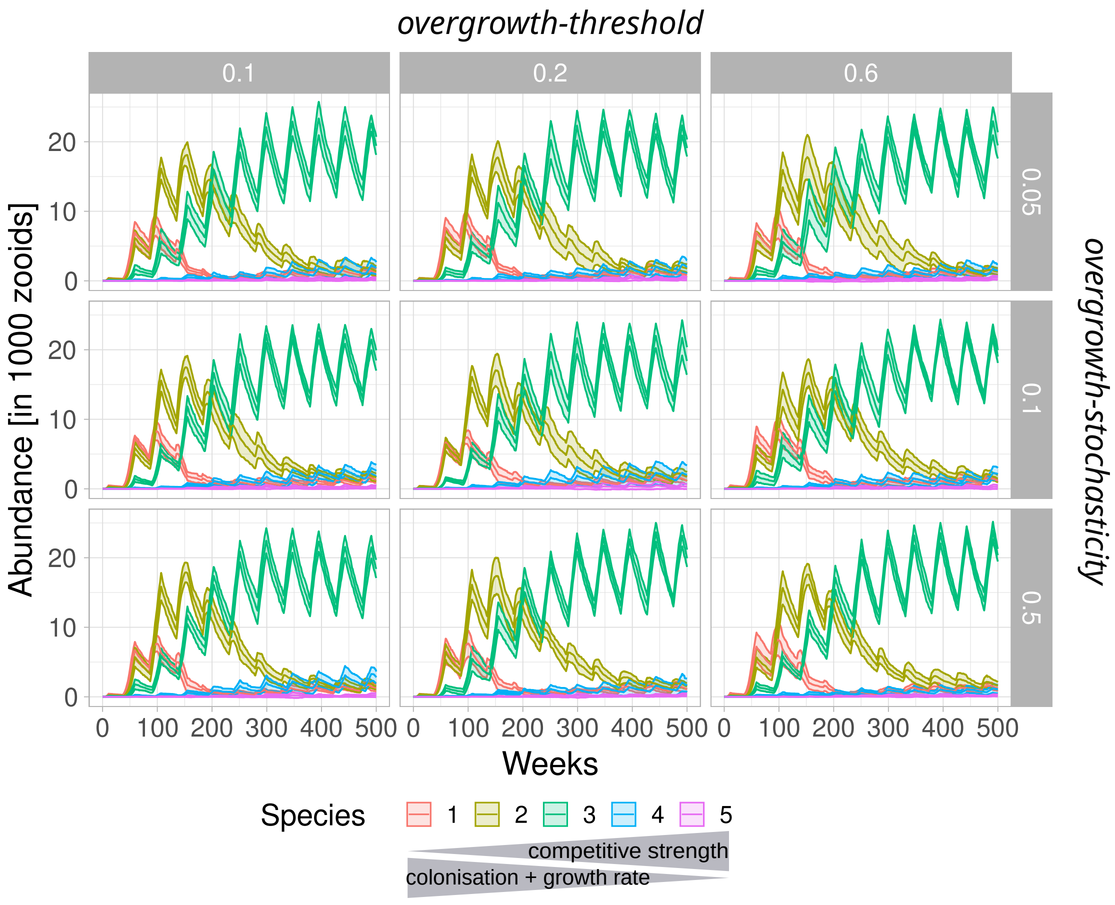


Supplementary Figure 4: **Successional dynamics with varying *overgrowth-threshold* and *overgrowth-stochasticity* parameters**. These two parameters influence the outcomes of overgrowths in our model. *overgrowth-threshold* determines the difference in competitive strengths necessary to produce clear overgrowth instead of tied outcomes. *overgrowth-stochasticity* determines the frequency of competitive reversals, where species loose overgrowth interactions despite having higher competitive ability. Both parameters were included based on empirical observations of overgrowth competition (see Methods for details). Here, we show that varying these parameter values does not affect the successional dynamics in our simulations and thus the results of our study. All simulations shown were run with a short season length and starting month was set to February. Data shown is the mean of 10 replicates, shaded areas correspond to standard deviation.
